# Systems-level analysis of transcriptome reorganization during liver regeneration

**DOI:** 10.1101/2021.10.18.464759

**Authors:** Manisri Porukala, P K Vinod

## Abstract

Tissue homeostasis and regeneration depend on the reversible transitions between quiescence (G0) and proliferation. The liver has a remarkable capacity to regenerate after injury or resection by cell growth and division. During regeneration, the liver needs to maintain the essential metabolic tasks and meet the metabolic requirements for hepatocyte growth and division. Understanding the regulatory mechanisms involved in balancing the liver function and proliferation demand after injury or resection is crucial. In this study, we analyzed high-resolution temporal RNA sequencing data of liver regeneration after two-thirds partial hepatectomy (PHx) using network inference and mathematical modeling approaches. The reconstruction of the dynamic regulatory network of liver regeneration reveals the trajectories of different metabolic pathways, protein processing in the endoplasmic reticulum (ER), ribosome biogenesis, RNA transport, spliceosome, immune response, and cell cycle. We further developed a mathematical model of the integrated circuit of liver regeneration that accounts for underlying features of compensatory metabolism, proliferation, and epithelial-to-mesenchymal transition during liver regeneration. We show that a mutually exclusive behavior emerges due to the bistable inactivation of HNF4A, which controls the initiation and termination of liver regeneration and different population-level expressions observed in single-cell RNA sequencing data of liver regeneration.

## Introduction

The liver is bestowed with an impeccable capacity to restore its lost mass following an injury or partial resection by coordinated cell growth (hypertrophy) and proliferation (hyperplasia). The ability to maintain and recover the original liver-to-body mass ratio is inferred as the thermostat-like regulator “Hepatostat”^1^. Different studies have used the surgical procedure of two-thirds partial hepatectomy (PHx) in rodents (Mus musculus, Rattus norvegicus) to understand liver regeneration ^2^. These studies have revealed the sources of regenerated liver mass and described the three phases in liver regeneration (priming, proliferation, and termination). The liver is the metabolically active organ and is at a crossroads of lipid and carbohydrates metabolism. The regenerating liver not only needs to maintain the essential metabolic function but also needs to meet the metabolic requirement of hepatocyte growth and division.

Liver regeneration depends on the control mechanisms regulating the reversible transition between quiescence and proliferation. Hepatocytes shift from quiescent to primed state with the expression of immediate-early (IE) genes in response to cytokines (IL6 and TNFα) derived from non-parenchymal cells^2-5^. The second phase of regeneration involves the activation of growth factor signalling. Non-parenchymal cells synthesize and release growth factors and promote the release of extracellular matrix (ECM)-bound reservoir of growth factors. These include growth factors HGF and EGF, which activate c-met and EGFR receptors, respectively^6, 7^. The last step involves cessation of proliferation by integrin signaling that promotes communication between ECM and epithelial cells^8, 9^. The liver-to-body mass ratio is maintained by controlling the rate of cell division and apoptosis. After two-thirds PHx, the hepatocytes also increase in size, followed by cellular division. An increase in the hepatocyte size alone is sufficient to recover the lost mass after PHx in Cdk1 knockout^10^. A significant decrease in NADH concentration and mitochondrial function is observed. This kind of compensatory mechanism is not without consequences since there is an increase in liver damage markers.

In addition to cytokines and growth factors, metabolic signals play a role in liver regeneration ^11, 12^. The change in metabolic demand under liver regeneration leads to systemic reorganization of metabolism. Animals subjected to PHx display hypoglycemia in the initial phase since the liver plays a major role in maintaining systemic glucose levels^13^. There is an increase in the systemic influx of lipids and triglycerides from extrahepatic adipose tissue after PHx leading to transient steatosis, which provides energy currency required for regeneration^12, 14, 15^. Other systemic cues include increased bile acid (BA) levels since the remnant liver cannot handle the BA returning via portal flow^16^. Blocking these metabolic alterations are shown to impair regeneration. Thus, regeneration is tightly intertwined with alterations in systemic metabolism.

Whole transcriptomic studies using microarray and RNA sequencing (RNA-seq) have mapped the gene expression pattern and transcriptional regulation in different liver regeneration models of rodents ^10, 17-25^. Metabolic genes are shown to be repressed, while the cell cycle genes are upregulated during liver regeneration. This raises the question of how the liver maintains metabolic homeostasis during liver regeneration. A division of metabolism into oxidative and biosynthesis phases has been proposed during liver regeneration ^26^. It is not clear how the liver achieves the dynamic balance between various cellular processes, including metabolism and cell cycle. Since the hepatocytes in the liver lobule are exposed to different microenvironments, there is a zonation (spatial heterogeneity) of gene expression. The liver lobule is metabolically partitioned to periportal, mid-lobular, and pericentral zones, with different zones exhibiting differences in proliferative capability ^27-29^. Recently, single-cell RNA sequencing (scRNA-seq) studies have started to reveal the division of labor with one population of hepatocytes activating early-postnatal-like gene expression and other compensating for metabolic function during liver regeneration ^30, 31^. In response to PHx, a wave of hepatocyte proliferation starting from zone 1 to zone 3 has been observed, with midzone 2 representing the primary source of new hepatocytes during liver homeostasis and regeneration ^32, 33^.

In this study, we modeled the temporal reorganization of the transcriptome of liver regeneration after PHx to understand the coordination of liver function and regeneration using a schema outlined in **Figure 1**. The inference of dynamic regulatory network from RNA-seq data was performed, which shows the interplay of different cellular processes at different time points during liver regeneration. The co-expression pattern of genes reveals the coordination of metabolism and the cell cycle. We also developed a mathematical model of the integrated circuit of liver regeneration, which accounts for the dynamic balance between requirements of liver function and regeneration as observed in scRNA-seq studies of liver regeneration.

**Figure 1:**
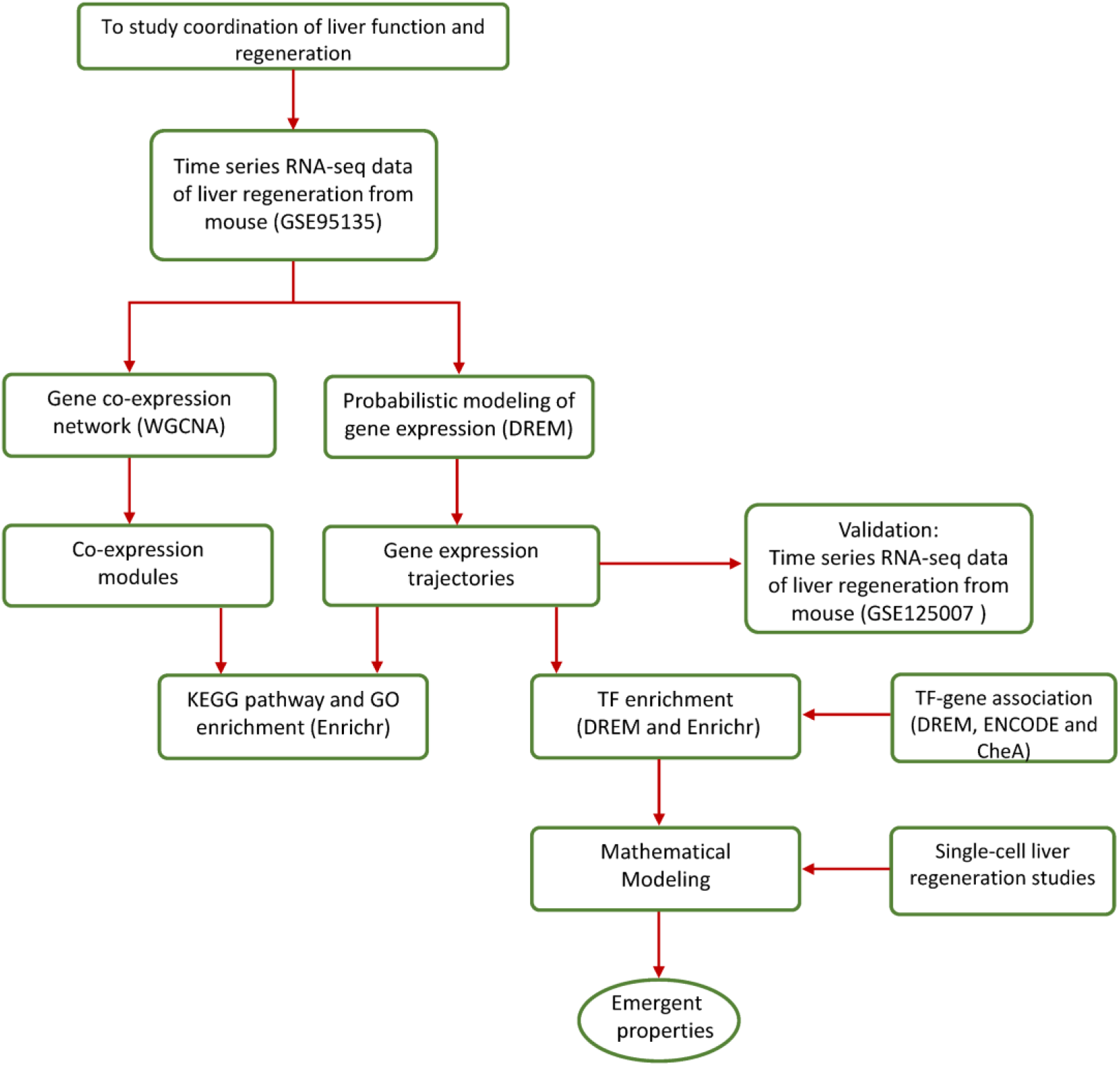
The workflow used to study the temporal reorganization of transcriptome during liver regeneration.

## Methods

### Transcriptomics data

We used the publicly available high-resolution temporal RNA-seq data (Illumina HISeq 2000) of liver regeneration after PHx from Gene Expression Omnibus (with accession number GSE95135)^23^. The samples in the dataset correspond to PHx operated 12-to 14-week-old male C57/BL6 mice entrained with 12 hours light-dark, fasting-feeding cycles. PHx samples include time points 0, 1, 4, 10, 20, 28, 36, 44, 48, 72, 168 and 672 hours. log_2_ +1 RPKM values were used for the downstream analysis. We also verified our findings using another RNA-seq data of liver regeneration (with accession number GSE125007)^24^. PHx samples include timepoints 0, 24, 30, 40, 48, 96, 168 and 672 hours.

### Reconstruction of co-expression network of liver regeneration

The co-expression network of liver regeneration was constructed using the weighted gene co-expression network (WGCNA) package in R ^34^. Top 5000 top varying genes across time points were selected (using rowVars function in R) to construct the correlation (Pearson) matrix for WGCNA. A linear transformation of correlation was performed to retain the sign of correlations.

A soft power adjacency function, a_ij_ = s_ij_^β^, was used to construct an adjacency matrix. We used the scale-free topology criterion to choose power β. This was obtained by computing the square of the correlation (R2) between log(p(k)) and log(k), where p(k) is the frequency distribution of the connectivity k. A plot of R2 and β, which shows a saturation characteristic, was used to choose the β value of 15. This corresponds to the point where the saturation is reached with mean connectivity ≥100. A topological overlap matrix (TOM) was constructed from the adjacency matrix, and 1-TOM was used to construct the dendrogram ^35^. The modules were identified using the dynamic tree cut algorithm with a minimum module size of 50. The module eigen gene (ME) expression was obtained by singular value decomposition (SVD). Enrichr was used to identify GO terms and KEGG pathways associated with each module ^36^.

### Probabilistic graphical modeling

We further reconstructed the dynamic regulatory network using the DREM method ^37^, which uses the Hidden Markov Model (HMM) to integrate time-series gene expression data with the transcription factor (TF)-gene association data. This approach clusters patterns of gene expression into paths and bifurcation points. Each bifurcation point represents a divergence in the expression of co-expressed genes under TF(s) influence. log_2_fold change in the expression of genes at every time point with respect to the reference time point (0 hours) was used as an input. We used the generated TF-gene association available for the mouse ^37^. The following parameters were used: a) Minimum log_2_ fold change of 1, b) The expression scaling weight set to 0.5, and c) TFs associated with a bifurcation point in the model were chosen with a hypergeometric distribution score less than 0.001. We also used CheA and ENCODE_and_CheA_consensus libraries ^36^ to identify significant (*adj p-value* < 0.05) TFs associated with the clusters.

### Mathematical Modeling

To model the regulatory circuit of liver regeneration, we adopted the framework proposed by Reinitz and colleagues ^38, 39^. This framework combines the best features of discrete and continuous approaches to simplify the complexity of the interactions in the network. We formulated a set of non-linear Ordinary differential equations (ODEs) of the form:

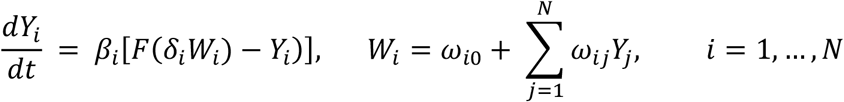

Y_i_ is expression level of the gene (0 ≤ *Y*_*i*_ ≤ 1), *F*(*δW*) = 1/(1 + *e*^−*δW*^) is “soft-Heaviside” function that varies from 0 (*W* << -1*/δ)* to 1 (*W* >>1*/δ*), *δ* determines the steepness of the function and *Wi* is the net effect on gene *i* of all genes in the network. The coefficient *ω*_*ij*_ can take values less than 0 (gene_*j*_ inhibits the gene_*i*_), more than 0 (gene_*j*_ activates gene_*i*_) or equal to 0 (no effect of gene_*j*_ on gene_*i*_). This equation also behaves like a discrete boolean for a large value of *δ*_*i*_’s. For *δ*_*i*_ values greater than 1, Y_i_ flips between 0 and 1 on a timescale ≈ β_i_^-1^. We studied how the qualitative behavior of the system changes with respect to parameter changes by performing phase plane and bifurcation analyses using XPPAUT (available from http://www.math.pitt.edu/~bard/xpp/xpp.html). The emergent properties of the liver regeneration network were analyzed.

## Results

### Dynamic regulatory network of liver regeneration

We first constructed the dynamic co-expression network of liver regeneration to study the transcriptome organization into functional modules using time-series expression data. We performed WGCNA and identified nine modules related to liver regeneration after PHx. Modules blue (M1), green (M2), red (M3), pink (M4), and purple (M5) show a positive correlation with pre-and post-PHx stages, while the modules black(M6), yellow (M7), brown (M8) and magenta(M9) show a negative correlation with stages (**Figure 2A**). However, the correlation of most modules with different time points of liver regeneration decreases. The eigen gene expression of each module shows the transient nature of gene expression with a change in direction occurring at different time points and recovery to the pre-PHx condition in the termination phase of liver regeneration (**Figure 2B and Figure S1**). The eigen gene expression of M4 and M5 modules increases between 1 and 4 hours, and of M2 and M3 modules increases between 4 and 10 hours post-PHx (priming phase). The eigen gene expression of the M1 module increases between 28 and 36 hours post-PHx (proliferative phase). The M5 module shows early recovery between 36 and 44 hours compared to other modules. The M8 and M9 modules are downregulated between 1 and 4 hours, while the M6 module is downregulated between 4 and 10 hours.

**Figure 2:**
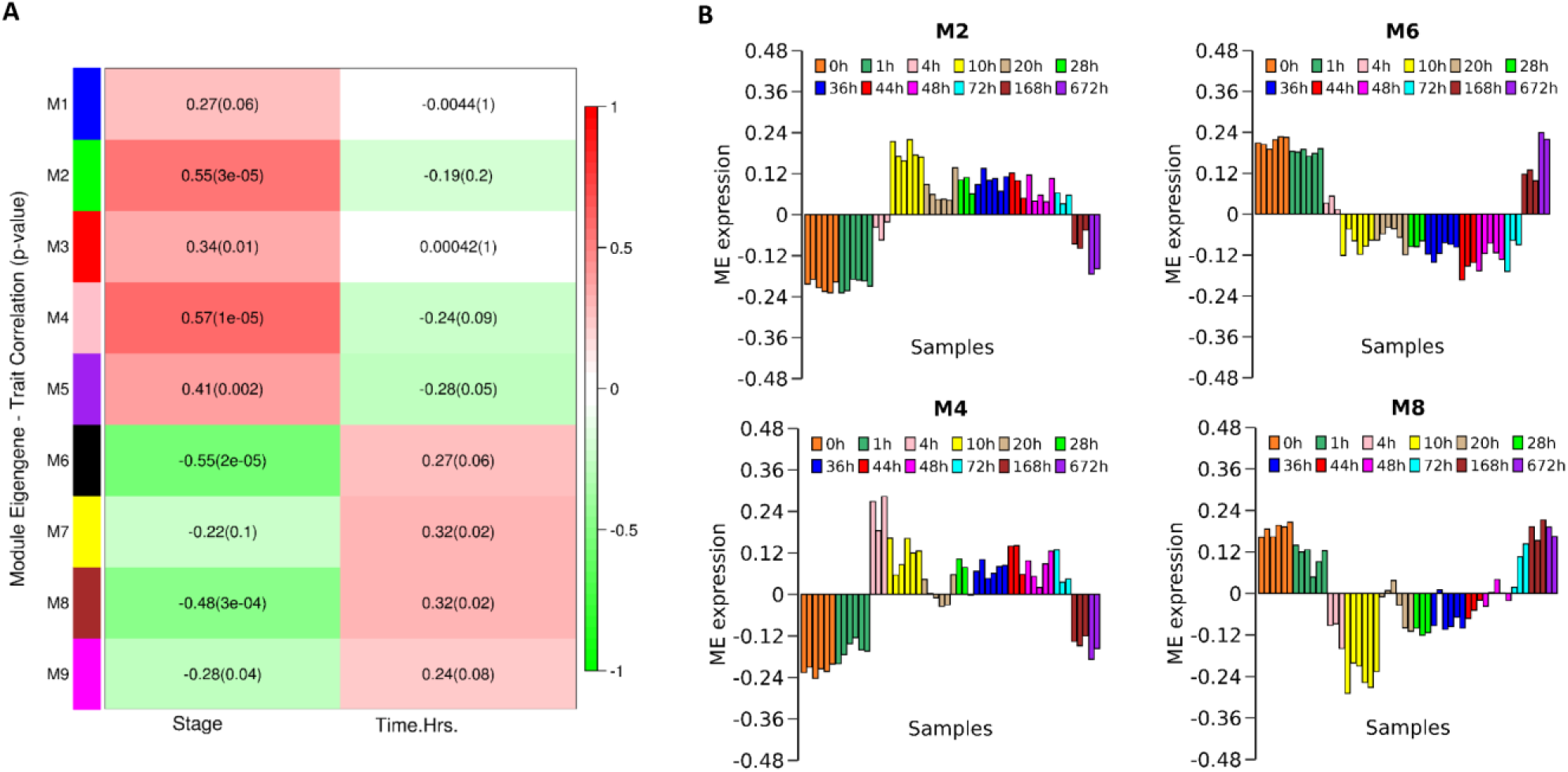
Modular organization of transcriptome of liver regeneration. **(A)** Correlation of module eigen (ME) gene expression with stages (pre-and post-PHx) and different time points. **(B)** Eigen gene expression profile of individual modules with respect to different time points of liver regeneration.

We also identified the biological processes and KEGG pathways associated with each module (**Table S1 and Data S1**). In the upregulated modules, the M1 module is associated with the cell cycle, DNA replication, and p53 signaling pathway; the M3 module is associated with protein processing in ER and protein export; the M2 is associated with complement and coagulation cascades, and the M4 module is associated with TNF signaling and ribosome biogenesis. The M1 module captures the proliferative response of hepatocytes that peaks after 36 hours, while the M3 module captures the role of endoplasmic reticulum (ER) stress. The upregulated modules also show a link with metabolic pathways: glutathione metabolism (M1), arginine and proline metabolism (M1), amino sugar, and nucleotide sugar metabolism (M3, M1), fatty acid degradation (M5 and M2), PPAR signaling pathway and peroxisome (M5).

The downregulated modules are primarily associated with metabolism (**Table S1**). The M6 module is associated with cholesterol metabolism, steroid hormone biosynthesis, bile acid biosynthesis, bile secretion, and PPAR signaling pathway. The M8 and M9 modules are associated with retinol and amino acid metabolism (branched-chain amino acids; glycine, serine, and threonine; tryptophan; cysteine and methionine; histidine). We also found glutathione metabolism, folate metabolism, pentose and glucuronate interconversions, Glycoxylate and dicarboxylate metabolism, and arachidonic acid metabolism as part of the downregulated modules. WGCNA revealed the global organization of liver transcriptome into modules, which are obtained based on the scale-free topology criteria.

To further generate insights into the dynamic organization and regulatory mechanism of liver regeneration, we performed probabilistic modeling of gene expression (see methods). This dynamic modeling approach revealed three core clusters that are upregulated immediately (cluster 1), upregulated after 28 hours (cluster 2), and downregulated immediately (cluster 3) post-PHx and their bifurcation into 17 sub-clusters (named paths A to Q) (**Figure 3 and Data S1**). We identified transcription factors (TFs) associated with the three core clusters. TFs regulating cluster 1 include FOS, JUN, CEBPB, NFKB, and STATs (**Table 1**). These TFs are known to be involved during the priming phase of liver regeneration. Other TFs include HNF4A, XBP1, LEF1, USF1, GATA4, EGR1, ESR1, and NFATCs. LEF1 is a downstream effector of the Wnt pathway important for hepatic periportal gene expression ^40^, and XBP1 is a regulator of unfolded protein response (UPR). Further, six sub-clusters (paths A to F) come under cluster 1 (**Figure 3A**). Paths A, B, C, and E are upregulated throughout the regeneration period up to 1-week post-PHx and return to the baseline at 4 weeks. Path A is enriched for complement and coagulation cascade, HIF and TNF signaling pathways (**Figure 4** and **Table S2**). Path B captures pathways related to protein processing in ER, amino sugar and nucleotide sugar metabolism, and protein export, which may play an essential role during the initial response soon after the resection. Path E is significantly upregulated post 10 hours and is enriched for fatty acid degradation, p53 signaling, and DNA replication. Path D shows an initial rise in gene expression, which returns to baseline 36 hours post-PHx (**Figure 3A**) and is enriched for metabolic pathways related to fatty acid and amino acid metabolism (**Figure 4**). Path F initially shows an increasing trend in gene expression but gets downregulated from 4 hours throughout the regeneration period, which makes it different from other paths in cluster 1. This path is enriched in steroid hormone biosynthesis and bile acid biosynthesis. The enrichment of paths D, E, and F suggests alterations in lipid metabolism in the priming phase.

**Table 1:**
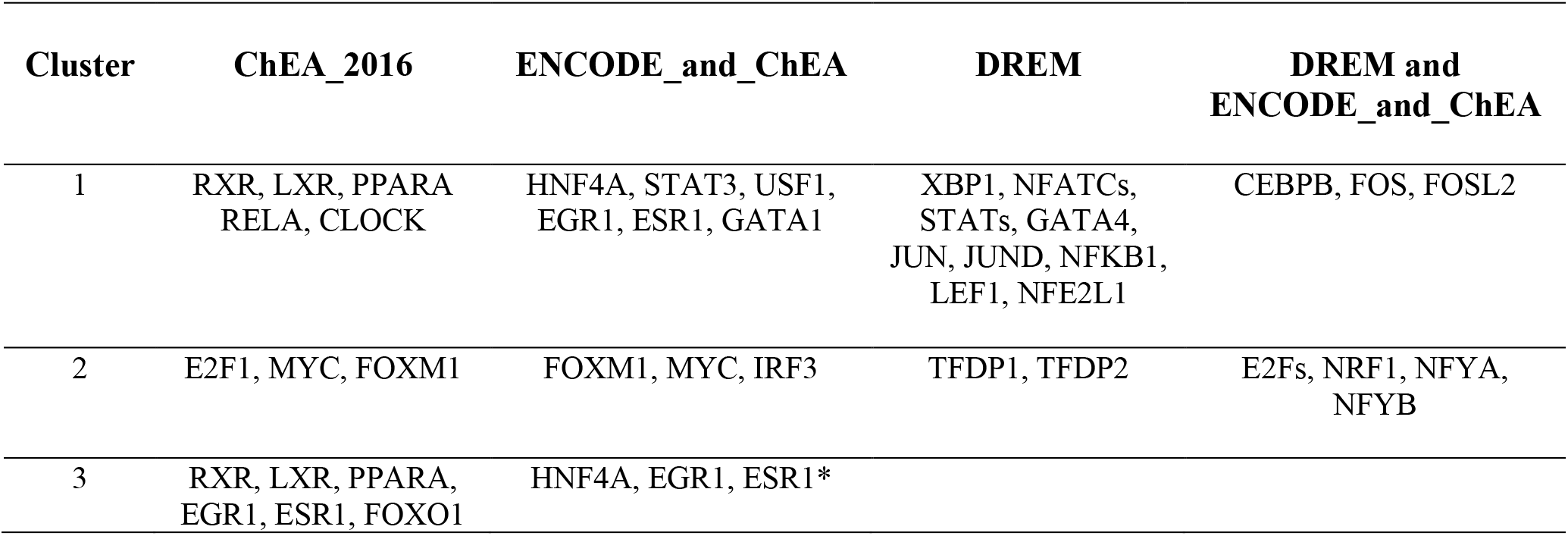
Transcription factors (TFs) associated with DREM core clusters. Significant TFs (adj p-value < 0.05) are identified based on different databases for transcription factor enrichment analysis. DREM results are based on generated TF-gene association for the mouse (score < 0.001). * represents uncorrected p-value.

**Figure 3:**
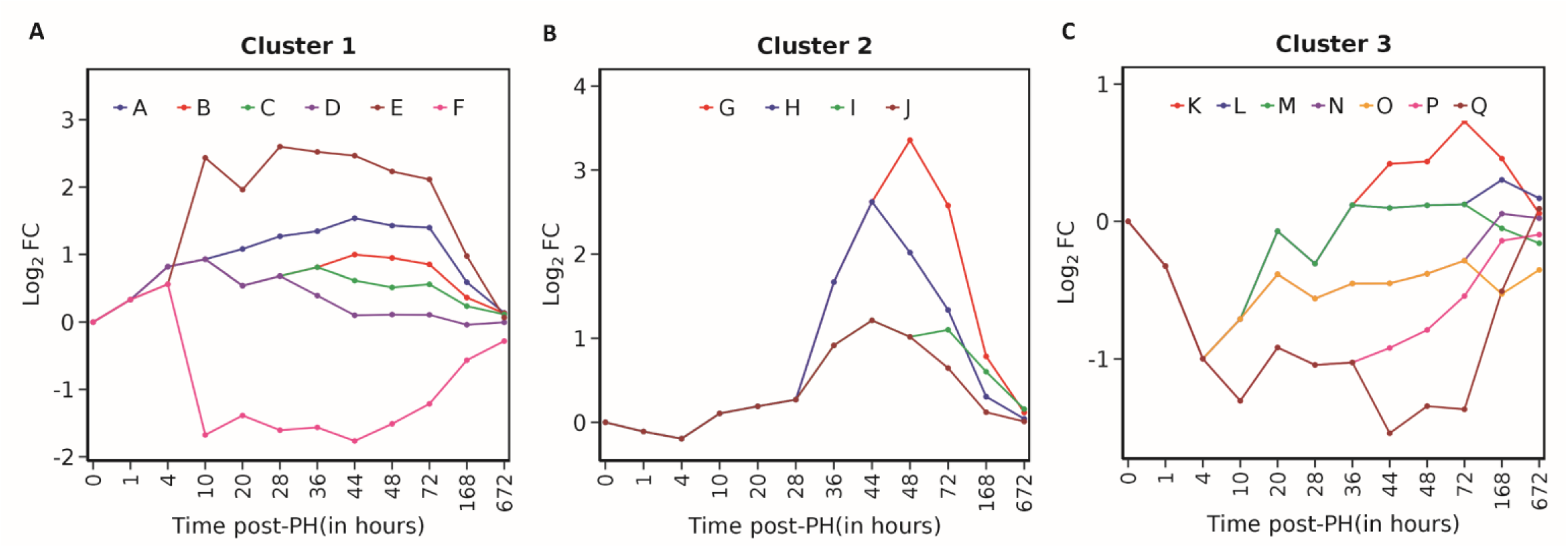
The regulatory paths of the set of co-expressed genes in the three core clusters of liver regeneration. The x-axis represents the time points of sample collection, and the y-axis represents the mean log2 fold change (log_2_FC) in mRNA expression post-PHx for each path. A path is split into multiple paths (split nodes) based on the divergence in gene expression.

**Figure 4:**
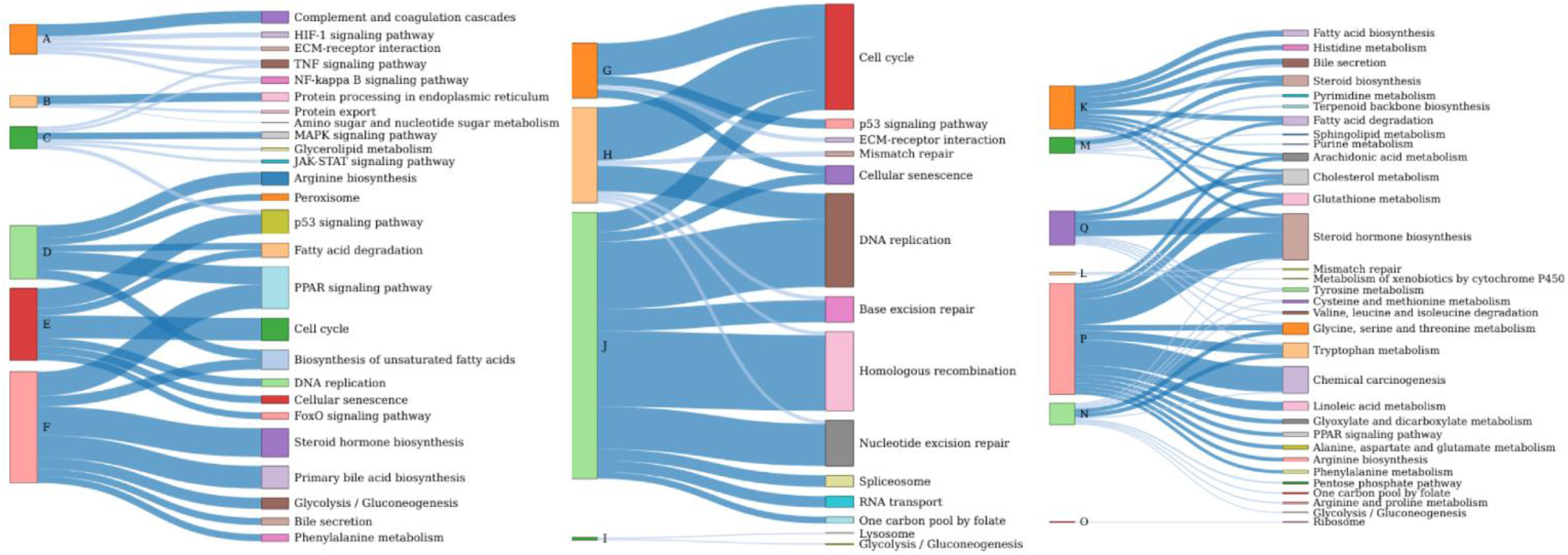
Sankey plot showing the significant KEGG pathways (dark blue: *adj p-value* < 0.05; light blue: *p-value* < 0.05; thickness represents -*log*_10_(*adi p*−*value*)) associated with different paths of cluster 1 (A to F), cluster 2 (G to J) and cluster 3 (K to Q) in the DREM analysis.

TFs regulating cluster 2 include E2Fs, FOXM1, MYC, NRF1, IRF3, NFYA, NYFB, and TFDP1/2 (**Table 1**). The gene expression of cluster 2 is relatively uniform compared to the other two core clusters. There are 4 paths (G to J) that come under cluster 2, showing an increasing trend post 28 hours (**Figure 3B**). Path G continues to increase until 48 hours and returns to baseline at 4 weeks. Paths G, H, and J of this cluster are significantly enriched for cell cycle events and DNA damage repair pathways (**Figure 4** and **Table S2**). Path I is associated with neutrophil degranulation. Cluster 3 shows both transient and sustained downregulation (paths K to Q) throughout the regeneration period returning to baseline after 1 week (**Figure 3C**). TFs regulating this cluster include HNF4A, RXR, LXR, EGR1, and ESR1 (**Table 1**). The paths of this cluster are mainly enriched for metabolic pathways (**Figure 4** and **Table S2**). Paths K, L, and M show transient downregulation at 4 hours; however, path K further rises during the proliferative phase. Path L is associated with negative regulation of JAK-STAT and cell size. Paths Q and P are downregulated throughout liver regeneration. Paths K, M, P, and Q are associated with the lipid and amino acid metabolism, with path K and P also associated with the glutathione metabolism. Paths N and O also show transient downregulation at 4 hours and are associated with amino acid metabolism and ribosome, respectively.

Further, we analyzed another recent RNA-seq data of liver regeneration using probabilistic modeling to confirm our findings. Although this dataset has a starting time point of 24 hours, we consistently observed the organization of the transcriptome into three core clusters pertaining to cell cycle, immune response, and metabolism with a similar set of transcriptional factors capturing the events of liver regeneration (**Figure S2** and **Table S3**). The changes related to the immune response are a continuum from the priming phase, as observed in **Figure 3A**. We also found pathways related to RNA transport and spliceosome upregulated in both datasets, while ER stress pathway is not enriched (**Table S3**). Lipid and glutathione metabolism are downregulated and co-cluster in a single path in both datasets (paths P and L in datasets 1 and 2, respectively). Downregulated steroid hormone biosynthesis and glycolysis/gluconeogenesis (cluster 3, paths L and N) are reset to the baseline by 96 hours. Cysteine and methionine metabolism and one-carbon folate pool are upregulated in the validation dataset.

### Alterations in liver metabolism during regeneration

The co-expression network-level analyses revealed the transient and sustained dynamical changes in metabolic processes post-PHx. We further analyzed the co-expression pattern of specific metabolic genes and pathways. Most genes of lipid metabolism are downregulated immediately, and their expression returns to the baseline between 36 and 72 hours, coinciding with the proliferative phase (**Figure S3 and S4**). We observed that genes related to the *de novo* lipogenesis pathway (SREBF1, FASN, ACACA, ACLY, ACSL3) and hydrolysis of fat (LPL) are further upregulated by 72 hours (cluster 3, path K) (**Figure S3**). HMGCR involved in converting HMG-CoA to mevalonic acid is initially downregulated along with SQLE (path K) (**Figure S5**). The mevalonate pathway plays an essential role in cholesterol synthesis. LDLR, which is involved in the uptake of cholesterol, is also downregulated initially. The initial phase of lipid metabolism downregulation coincides with the cell growth phase (hypertrophy before the initiation of proliferative phase) observed after PHx. A negative correlation between lipid metabolism and cell size has been reported ^41^. Further, the RNAi of SREBs that are involved in lipid metabolism results in cell size increase.

In the initial phase of regeneration, along with lipid metabolism, we also observed a decrease in mitochondrial gene expression (ATP5J2, NDUFA6, NDUFV3, PDK2) (**Figure S6**). However, the expression of some of the related genes (UCP2, PDK4 and COX6B2) increases during liver regeneration. UCP2 is involved in uncoupling substrate oxidation from ATP synthesis. It is a negative regulator of mitochondrial superoxide production that modulates cell proliferation ^42^. PDK4 is a key regulator coordinating lipid metabolism with liver growth ^43^. Mitochondrial metabolism is linked to mammalian cell growth and proliferation ^41^. The RNAi of PGC1α, which controls the expression of genes regulating oxidative phosphorylation, TCA cycle, and lipid synthesis, increases cell size. A decrease in mitochondrial gene expression in the initial phase of liver regeneration is accompanied by upregulation of LDHA (cluster 1, path A) involved in anaerobic glycolysis. On the other hand, high mitochondrial and lipogenic transcriptional programs are required to promote proliferation^44^.

Genes of glutathione metabolism are also transiently downregulated (cluster 3, paths P and L) (**Figure 5**). These include GCLC and GCLM involved in *de novo* synthesis of GSH, which plays an important role in scavenging reactive oxygen species and maintaining the redox balance. GCLC and GCLM encode the catalytic and modifier subunits of glutamate cysteine ligase (GCL), respectively. GCL catalyzes the rate-limiting step involved in the generation of γ-glutamylcysteine (γ-GC) from glutamate and cystine. γ-GC and GSH levels are known to be regulated by inflammatory signaling^45^. However, glutathione S-transferases (GSTM1, GSTM2, GSTM3, GSTM4, GSTM6, and GSTM7) are transiently downregulated in 10 hours. Increased GSH levels are reported in HCC and during liver regeneration ^46^. GSH deficiency interferes with liver regeneration after PHx ^47^. In the NAD salvage pathway, we observed that NNMT is upregulated (cluster 1, path E) during liver regeneration while NAMPT levels fluctuate with downregulation at 4 and 28 hours (cluster 3, path L) (**Figure 5**). However, NAMPT is upregulated at 36 hours. Deletion of NAMPT is shown to affect cell proliferation during liver regeneration^48^. NNMT knockdown leads to an increase in lipogenic gene expression and a decrease in gluconeogenic gene expression. Both NAMPT and NNMT control lipid, cholesterol, and glucose metabolism by stabilizing SIRTs ^49^. NNMT is at the crossroad of metabolism and epigenetic regulation, but in the liver, it is not the major methyltransferase to maintain S-adenosyl-methionine (SAM) to S-Adenosyl homocysteine (SAH) ratio^50^. However, NNMT overexpression can decrease NAD levels, reduce methylation capacity, and promote liver steatosis and fibrosis^51^.

**Figure 5:**
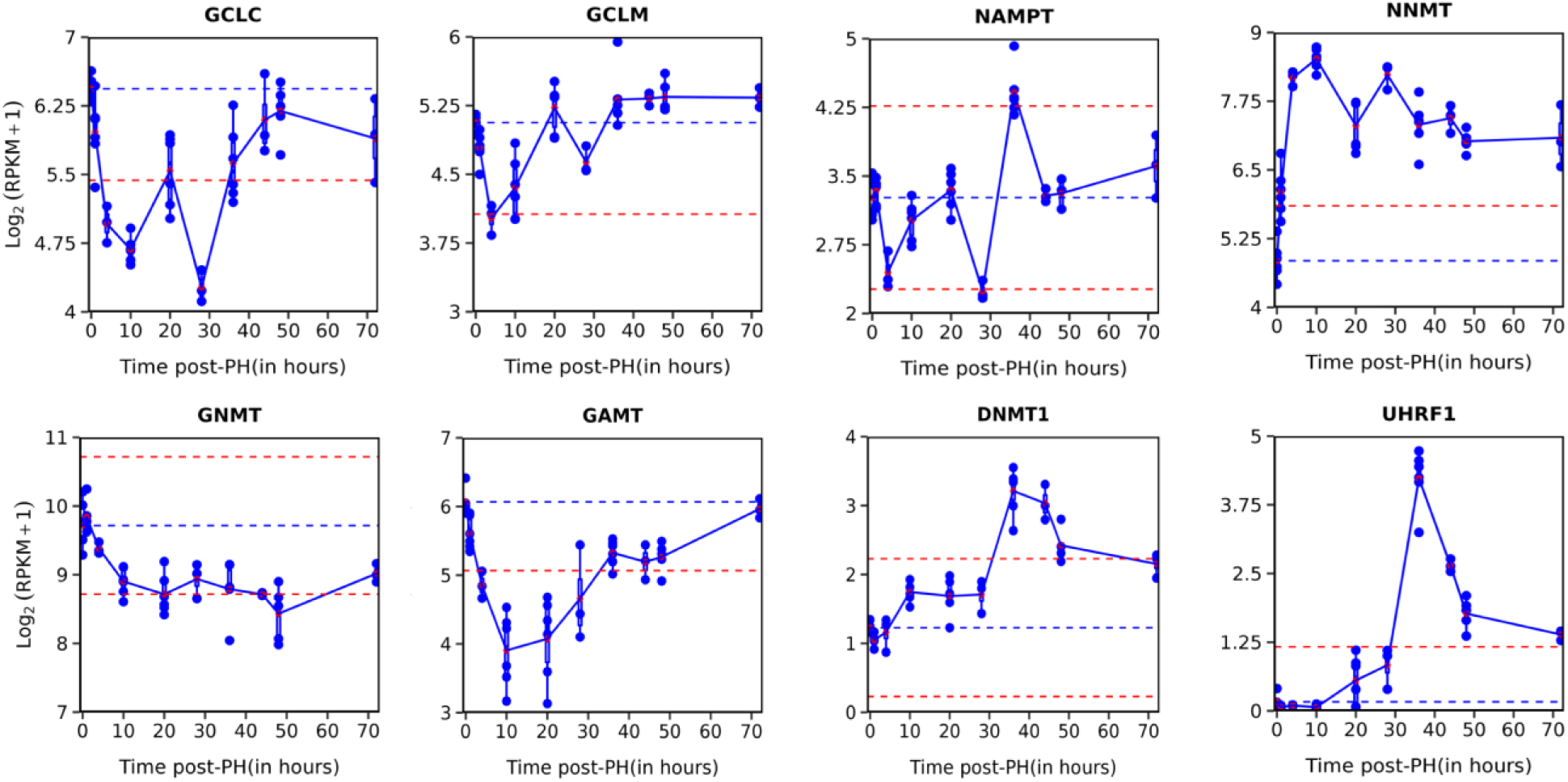
Expression profile of genes affecting GSH, NAD, and SAM levels. The blue line represents the baseline (0 hours), and the red line represents a two-fold change with respect to the baseline.

We observed that the expression of main methyltransferases of the liver (GNMT, GAMT) involved in the conversion of SAM to SAH is downregulated (cluster 3, path P), and genes of methionine catabolism (MAT1A and MAT2A) involved in the production of SAM are upregulated (cluster 1, paths B and E). This shows that SAM levels may increase during regeneration and contribute to epigenetic control. This is further supported by the upregulation of genes involved in DNA methylation (DNMT1 and UHRF1) (cluster 2, path H) (**Figure 5**), which is consistent with the observation by Wang *et al*., (2019). Both these genes are co-expressed with cell cycle genes in the mid-phase, while genes involved in SAM production are upregulated very early on. On the other hand, we observed that genes of different amino acids metabolism (BCAA, Tryptophan, and Tyrosine metabolism) are also transiently downregulated. A decrease in expression of genes of the BCAA degradation pathway is consistent with observations that the levels of BCAA in plasma increase during liver regeneration ^11, 12^.

### Alterations in liver zonation

Hepatocytes segregate into three functional zones (perivenous, mid-zonal, and periportal hepatocytes), referred to as liver zonation. Halpern and colleagues have reported gene expression specific to these regions ^27^. We examined how zone-specific marker genes are affected by PHx. Periportal genes ARG1, ASL are upregulated, while ASS1, ALB, and CYP2F2 are downregulated (**Figure S7**). The expression of periportal-specific transcriptional factor YAP1 shows an increasing trend along with its targets CTGF and CYR61. The pericentral gene CYP2E1 is significantly downregulated, while HIF1A, AXIN2, OAT, and ANG are upregulated (**Figure S8**). The mid-zonal gene HAMP is upregulated, while MUP3 and CYP8B1 are downregulated (**Figure S9**). Although zone-specific gene expression shows a mixed pattern including compensation with the increase in the expression of some of the genes, our results indicate that metabolic functions such as cholesterol biosynthesis, bile biosynthesis, and lipogenesis are transiently downregulated. These results suggest that a transient decrease in hepatic metabolism may counterbalance hepatocyte function vs. hepatocyte growth and proliferation.

### A model of balance between liver identity and proliferation during liver regeneration

The transcription factor enrichment analysis showed HNF4A as a TF governing cluster 1 and cluster 3 (**Table 1**). Liver identity genes (223 out of 622 identified genes)^52^ show overlap with these two core clusters that are upregulated and downregulated immediately. On the other hand, there is no overlap with cluster 2, the upregulated cell cycle cluster activated post 28 hours. We also observed that there is less overlap between liver identity genes and cell cycle clusters in the validation dataset. This pattern of gene expression may be due to the segregation of functions among hepatocytes. However, the immediately upregulated cluster also includes some genes of cell cycle and p53 signaling pathway activated early compared to the cell cycle cluster activated post 28 hours. These include Cyclin D (CCND1) and Cdk inhibitor (CDKN1A); their ratio controls the passage through the restriction point and G1/S transition (**Figure S10**). Interestingly, AFP that is induced in the fetal liver is also co-expressed with cell cycle genes activated post 28 hours (path J).

Recently, single-cell transcriptomic data of liver regeneration has shown that there is a functional bifurcation of hepatocytes into proliferative and non-proliferative cells^30^. The hepatic function genes are upregulated predominately in non-proliferating cells, while these genes are downregulated in dividing cells. This suggests that there is a division of labor. The staining for Hnf6 (Onecut1) and Hnf4a, two key hepatocyte transcription factors, decreased in replicating cells^53^. A mathematical model of a simple regulatory circuit was developed to demonstrate how the division of labor occurs. We propose that a mutually exclusive behavior between liver function and cell division can be established by the feedback loop regulation between these processes during liver regeneration. The regulatory circuit includes mutual antagonism between Cyclin D and HNF4A ^54-57^ (**Figure 6A**). Deletion of HNF4A results in increased expression of Myc and Cyclin D, while Cyclin D represses the transcriptional activity of HNF4A. This double negative feedback loop circuit can be regulated by a plethora of signals (local and in the circulation) activated during the priming phase of liver regeneration. The PHx can alter the balance between mitogen and mitoinhibitors by matrix remodeling, induce secretion of ligands and cytokines and change the circulating levels of metabolites in plasma. Further, the underlying mechanism of control of this core double negative circuit can vary since hepatocytes express different genes depending on their location in the hepatic lobule along the periportal-pericentral axis. We propose two kinds of inputs: proliferative and compensation signals that can act on the core circuit in a context-dependent manner in different hepatocytes to bring about different outcomes. The proliferative input suppresses HNF4A and activates Cyclin D expression, while compensation signals promote HNF4 activation.

**Figure 6:**
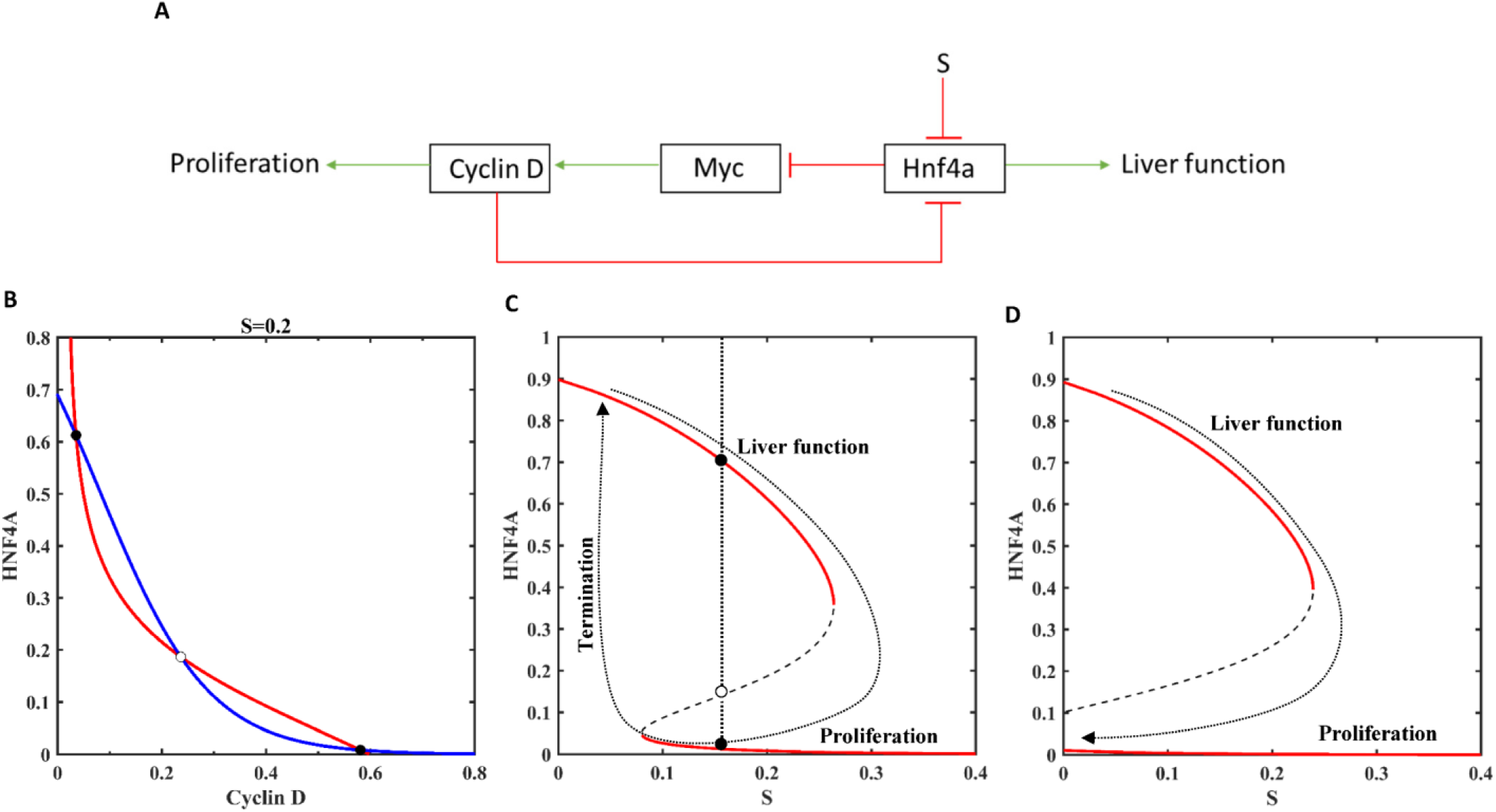
A model of balance between liver function and regeneration. **(A)** A core regulatory circuit of mutual antagonism between Cyclin D and Hnf4a controlled by inhibitory input stimuli S is shown. The activation is shown in green and inhibition in red. **(B)** Phase plane showing the nullclines of Hnf4a (blue) and Cyclin D (red) for S=0.2. **(C)** Bistable inactivation of Hnf4a with an increase in input stimuli S. **(D)** Irreversible inactivation of Hnf4a with an increase in feedback loop strength (w_HNF4A_CycD_= -1.5). The solid circle represents the stable steady state, and the open circle represents the unstable steady state.

The phase plane analysis of the model shows that the nullclines of HNF4A and Cyclin D intersect at three points creating two stable and one unstable steady states (**Figure 6B**) for the parameter values given in **Table S4**. This circuit exhibits bistable characteristics depending on the strength of the proliferation signal S (**Figure 6C**). The two stable states correspond to hepatocytes in replicating (high Cyclin D) and differentiated (high HNF4A) states. At the intermediate signal strength, two populations of hepatocytes (differentiated and replicating state) co-exist. The liver regeneration program after PHx can be viewed as changes occurring around the bistable switch. This leads to transient loss of hepatocyte identity that facilitates the process of regeneration. The re-activation of HNF4A with the decrease in the input signal (due to repair) becomes essential for the termination of liver regeneration (shown by arrow in **Figure 6C**). The development of HCC can be explained by the shift in re-activation threshold to a negative regime with the change in feedback loop strength, making the transition irreversible (**Figure 6D**). On the other hand, the compensation signal shifts the HNF4A nullcline up resulting in one stable steady state corresponding to hyperactivation of HNF4A as observed by Chembazhi *et al*., (2021) (**Figure 7**).

**Figure 7:**
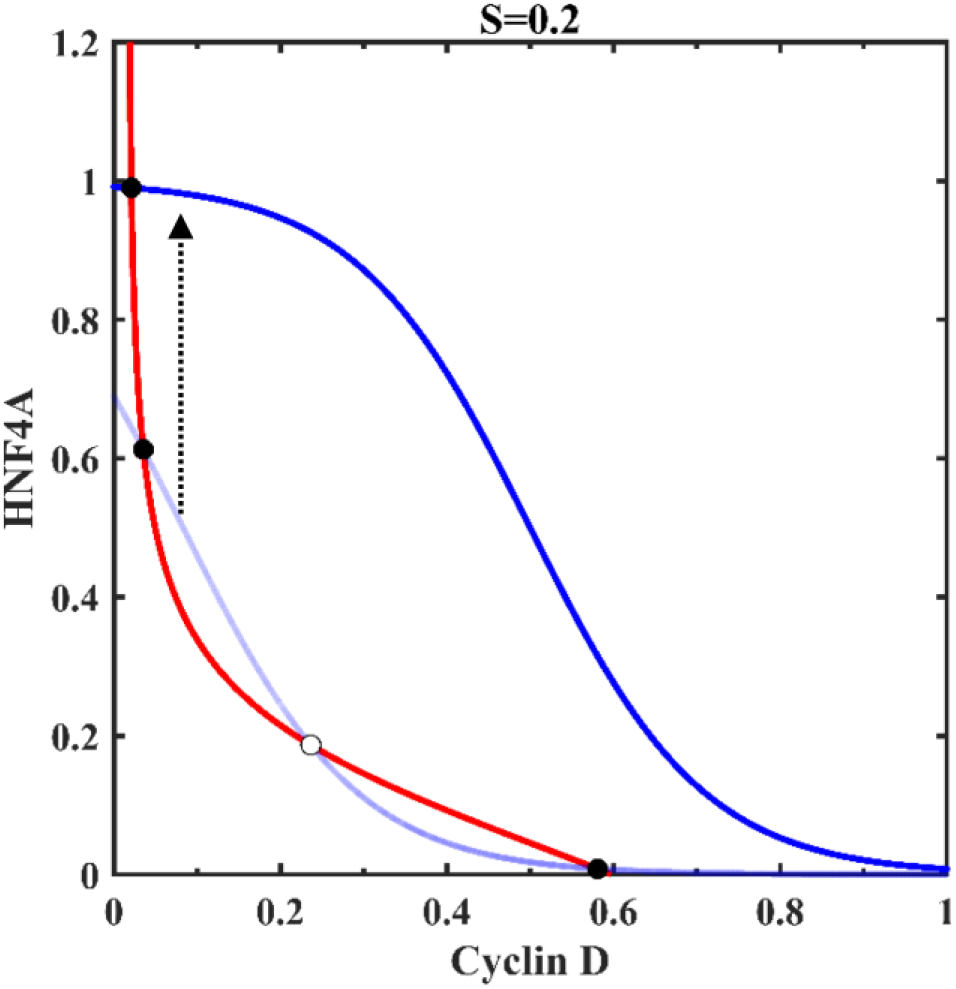
Phase plane showing the hyperactivation of Hnf4a during liver regeneration. The nullcline of Hnf4a (blue) shifts above (arrow) in the presence of activatory stimuli (M=0.5). Nullclines of Cyclin D (red) and Hnf4a (light blue) for inhibitory stimuli S=0.2 are given for reference. The solid circle represents the stable steady state, and the open circle represents the unstable steady state.

Further, the repression of the mesenchymal program is also required for the maintenance of liver identity ^58^. HNF4A and epithelial-to-mesenchymal transition (EMT) master regulatory gene SNAIL form a mutual inhibition circuit, which controls the balance between liver differentiation and mesenchymal program (**Figure 8A**). EMT control network involves SNAIL-induced self-activation of ZEB1^59^. An inhibition of HNF4A by proliferative signal can activate EMT in some hepatocytes. The phase plane analysis of this control circuit shows tristability (co-existence of three stable steady states) depending on the strength of proliferation signal S (**Figure 8B to D**) for the parameter values given in **Table S4**. These states correspond to (1) high HNF4A with low SNAIL/ZEB1, (2) low HNF4A with high SNAIL/ZEB1, and (3) high HNF4 and SNAIL/ZEB1 (hybrid state). The single-cell omics study of liver regeneration has shown the existence of a hybrid cluster enriched for epithelial and hepatocyte-specific features and markers of mesenchymal cells ^30^. The proposed regulatory circuit accounts for the cellular plasticity during liver regeneration.

**Figure 8:**
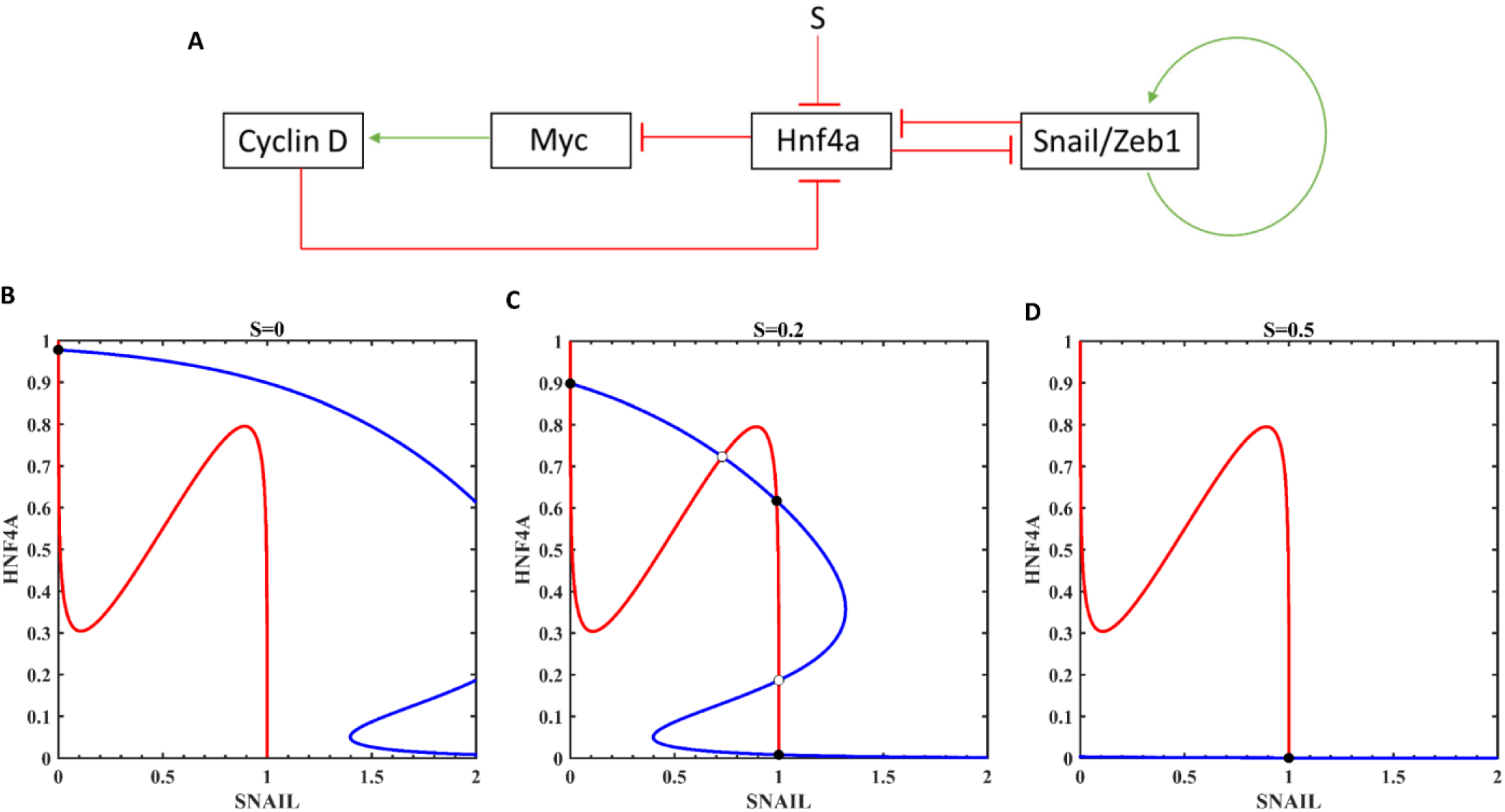
Multistability of the underlying circuit of liver regeneration. **(A)** The regulatory circuit controlling the balance between liver function and EMT during liver regeneration is shown. Activation is shown in green and inhibition in red. Nullclines of Hnf4a (blue) and Snail/Zeb1(red) are shown for different input stimuli **(B**) S=0, **(C)** S=0.2, and **(D)** S=0.5. The solid circle represents the stable steady state, and the open circle represents the unstable steady state.

## Discussion

The liver balances its function and proliferation demand after injury or resection. Recent advancement in high throughput techniques is helping to understand further the regulatory mechanisms involved in the regulation of liver regeneration. We analyzed RNA-seq data of liver regeneration to understand the temporal reorganization of the transcriptome and to generate network-level insights. We also analyzed the results obtained based on scRNA-seq of liver regeneration. The dynamic network reconstruction revealed the trajectory of major pathways that are upregulated and downregulated (transient vs. sustained) during liver regeneration (**Figure 2 and 3**). Our analysis supports the model of mutual antagonism between liver function and proliferation in liver regeneration. We show multistability of the underlying network of liver regeneration.

While overall metabolic downregulation suggests a decrease in liver function, the dynamics of metabolic pathways suggest that maintaining the levels of specific metabolites are required for liver regeneration. We observed that fine-tuning of SAM levels might be required for the methyltransferase reactions in liver regeneration (**Figure 5**). This is supported further by the downregulation of major liver methyltransferases. The co-expression pattern of cell cycle and DNA methylation genes highlights the scenario for crosstalk between cell cycle and chromatin regulatory proteins. Genes of NAD, glutathione, and lipid metabolic pathway decreased and reappeared immediately in the priming phase of liver regeneration. This suggests a possible requirement of these pathways for the cell cycle progression. Although NNMT expression correlates with adiposity, its expression during liver regeneration may be beneficial. Hepatic steatosis is shown to alter the demand for NAD and GSH ^60^. Lipid metabolic genes are also further upregulated at 36 to 72 hours, respectively, coinciding with the proliferative phase (**Figure S3**). On the other hand, PPARA, a transcriptional activator that controls β-oxidation, follows the inverse profile with its expression returning to baseline during the proliferative phase. Along with a decrease in the liver’s metabolic function, our analysis also captures the dynamic changes in metabolism that may indicate the shift from growth to proliferative phase during liver regeneration.

Another pathway that is upregulated during liver regeneration is protein processing in ER and protein transport (**Figure 4 and Table S1**). ER stress plays a role in liver metabolism, damage, and inflammation^61^. The knockdown of XBP1 results in liver injury and impairment of liver regeneration^62^. Loss of IRE1, an upstream activator of XBP1, impairs liver regeneration with activation of STAT3 affected^63^. ER stress is also shown to suppress the liver identity gene in the damaged liver^52^. On the other hand, an increase in ER stress under HFD conditions can impair liver regeneration^64^. We also found ribosome biogenesis and RNA processing as important features of liver regeneration. Ribosome biogenesis is known to increase during cell growth and proliferation^65^. PTBP1 involved in the regulation of alternate splicing events is co-expressed with the cell cycle cluster 2. On the other hand, genes involved in alternate polyadenylation (SRSF3 and SRSF7 are co-expressed) of mRNA precursors^66^ are immediately upregulated post-PHx compared to PTBP1. A decrease in SRSF3 expression is observed in mouse models of NAFLD and NASH ^67^. A global change in alternate splicing machinery is observed in NAFLD^68^. These results suggest that mRNA cleavage and polyadenylation may also control gene expression during liver regeneration in addition to epigenetic regulation. Consistently, 3,5 diethoxicarbonyl-1,4 dihydrocollidine (DDC) treatment leads to liver regeneration and a switch to a fetal splicing program ^22^.

The co-expression pattern of genes suggests a mutually exclusive behavior of the cell cycle and liver identity genes during liver regeneration. Few liver identity genes were upregulated and were co-expressed with Cdk inhibitor (CDKN1A) and activator (CCND1). An increase in Cdk inhibitor level may provide a window of opportunity for hepatocytes to grow before dividing. Alternatively, the co-expression of liver identity genes with Cdk inhibitor may suppress the cell division and maintain liver function^69^. On the other hand, Cyclin D expression alone influences the transcriptional regulation of liver metabolism^55^. Distinguishing these effects requires single-cell level quantification. Mathematical modeling showed that interaction between regulators of cell cycle and liver function could make the system bistable (**Figure 6**), which accounts for the co-existence of two populations of hepatocytes, with one undergoing cell division while the other helping to maintain liver function^30, 31^. The bistable switch accounts for transient inactivation of HNF4A with dynamic change in input signals during liver regeneration. We highlighted that the transition from liver function to proliferation could become irreversible with a change in the feedback loop strength. In this picture, the termination of liver regeneration depends on the reactivation of HNF4A, which is consistent with ^70^. Different studies have reported HNF4A inactivation in HCC^71-73^. We also showed that multistability emerges by coupling the HNF4A feedback loop with the EMT circuit (**Figure 8**). The EMT circuit is also known to exhibit tristability in cancer progression ^74^.

We propose an integrated circuit of liver regeneration by extending the core circuit (**Figure 9**). The cell cycle and EMT control during liver regeneration may involve the YAP1/Hippo and Wnt/β-catenin signaling pathways converging on HNF4A inactivation. Yap1, a mechanical rheostat acting downstream of the Hippo signaling pathway, has a direct role in hepatocyte differentiation by inhibiting HNF4A and activating SNAIL^75^. Reciprocal control of Yap1 by SNAIL (activating) and HNF4A (inhibiting) has been shown, resulting in a complex circuit of multiple feedback loops controlling liver identity. The ectopic activation of Yap1 is sufficient to de-differentiate hepatocytes into cells with stem cell-like characteristics^76^. The early phase of liver regeneration is accompanied by Yap1 activation and nuclear location^77^. Yap1 also cooperates with Myc in the control of proliferation^78^. This mechanism explains the existence of proliferative hepatocytes undergoing EMT during liver regeneration^30^.

**Figure 9:**
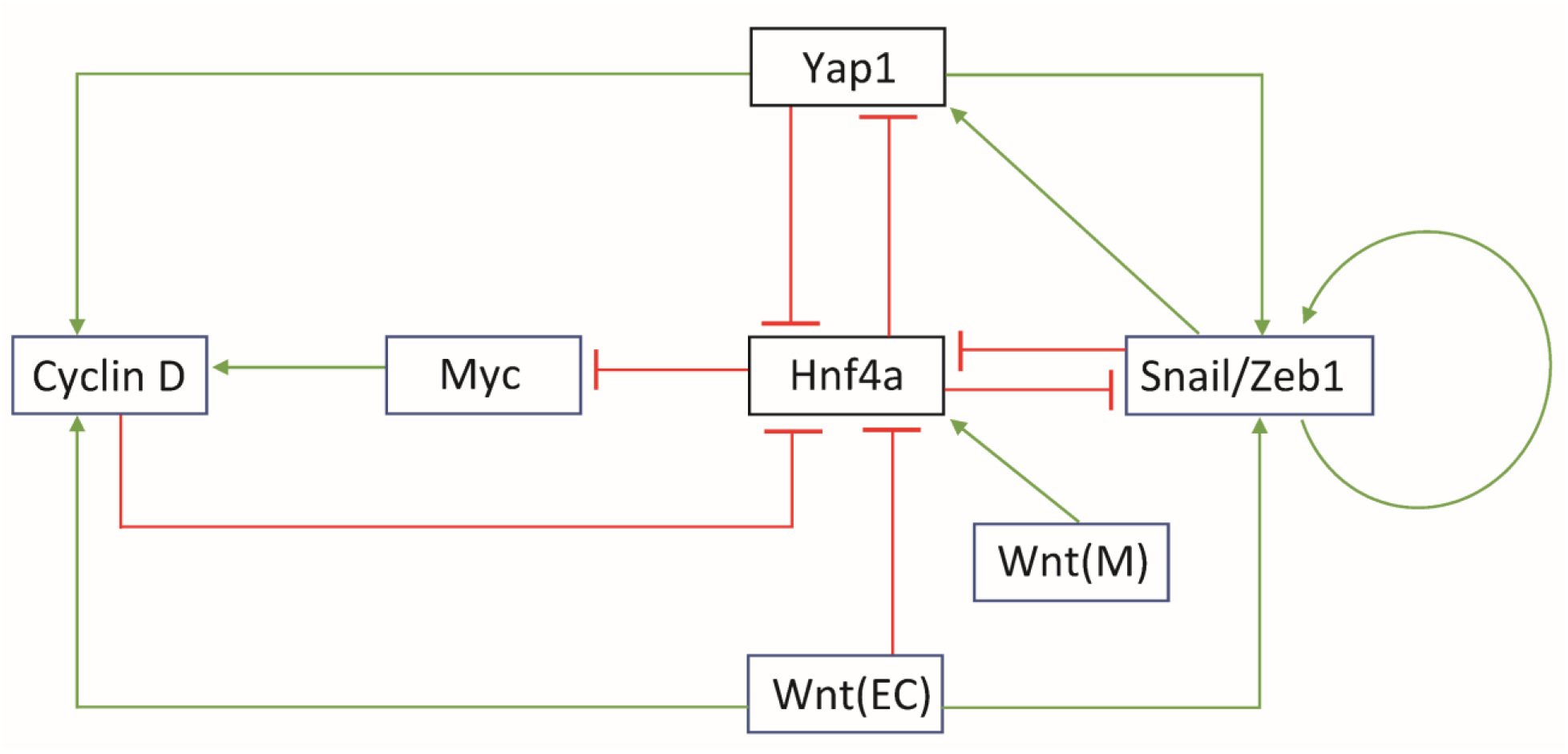
The proposed integrated circuit of liver regeneration controlled by Yap1 and Wnt. Activation is shown in green and inhibition in red. EC-endothelial cell, M-macrophage.

Wnt/β-catenin signaling pathway is also induced in response to liver regeneration under PHx ^79, 80^. Wnt can also control the same circuit of HNF4A and EMT. It is known that Wnt and HNF4A mutually inhibit each other, and Wnt activates SNAIL to control the EMT^81^. Myc and β-catenin cooperate in liver carcinogenesis with Yap1 as a mediator^82^. It is also shown that sinusoidal endothelial cell Wnts drive proliferation, while macrophage Wnts drive functional compensation ^69^. This integrated circuit of liver regeneration controlled by Yap1 and Wnt may provide underlying features of proliferation, compensatory metabolism, and EMT states as observed in single-cell studies. Thus, simultaneous control of HNF4A may drive the bifurcation of hepatocytes into different activity states. On the other hand, both Wnt and Yap1 have opposing functions to establish liver zonation ^79^. Wnt/β-catenin signaling is active in pericentral hepatocytes, while YAP1 is expressed in the periportal region. It will be interesting to study further the factors that control the dual role of Yap1 and Wnt in liver regeneration and zonation. Overall, our study provides a systems-level view of liver regeneration post-PHx. The underlying gene modules identified here can be connected to phenomenological model of liver regeneration ^83^ to obtain the dynamical characteristics of entry and exit from liver regeneration.

## Author Contributions

Conceptualization, P.K.V.; Data curation and analysis, M.P.; Investigation, M.P. and P.K.V.; Funding acquisition, P.K.V.; Writing-original draft, M.P. and P.K.V.; All authors have read and agreed to the published version of the manuscript.

## Data Availability Statement

All the datasets are freely available and can be downloaded from https://www.ncbi.nlm.nih.gov/geo/ using the given accession numbers.

## References

1. G. K. Michalopoulos, The American journal of pathology, 2010, 176, 2–13.

2. G. K. Michalopoulos and B. Bhushan, Nat Rev Gastroenterol Hepatol, 2021, 18, 40–55.

3. A. I. Su, L. G. Guidotti, J. P. Pezacki, F. V. Chisari and P. G. Schultz, Proceedings of the National Academy of Sciences, 2002, 99, 11181–11186.

4. R. Taub, Nature reviews Molecular cell biology, 2004, 5, 836–847.

5. N. Fausto, J. S. Campbell and K. J. Riehle, Hepatology, 2006, 43, S45–S53.

6. C.-G. Huh, V. M. Factor, A. Sánchez, K. Uchida, E. A. Conner and S. S. Thorgeirsson, Proceedings of the National Academy of Sciences, 2004, 101, 4477–4482.

7. A. Natarajan, B. Wagner and M. Sibilia, Proceedings of the National Academy of Sciences of the United States of America, 2007, 104, 17081–17086.

8. V. Gkretsi, U. Apte, W. M. Mars, W. C. Bowen, J. H. Luo, Y. Yang, Y. P. Yu, A. Orr, R. St-Arnaud, S. Dedhar, K. H. Kaestner, C. Wu and G. K. Michalopoulos, Hepatology, 2008, 48, 1932–1941.

9. S. Donthamsetty, W. Bowen, W. Mars, V. Bhave, J. H. Luo, C. Wu, J. Hurd, A. Orr, A. Bell and G. Michalopoulos, Toxicol Sci, 2010, 113, 358–366.

10. M. J. Caldez, N. Van Hul, H. W. L. Koh, X. Q. Teo, J. J. Fan, P. Y. Tan, M. R. Dewhurst, P. G. Too, S. Z. A. Talib, B. E. Chiang, W. Stunkel, H. Yu, P. Lee, T. Fuhrer, H. Choi, M. Bjorklund and P. Kaldis, Developmental cell, 2018, 47, 425–438 e425.

11. J. Huang and D. A. Rudnick, The American journal of pathology, 2014, 184, 309–321.

12. D. A. Rudnick and N. O. Davidson, International journal of hepatology, 2012, 2012, 549241.

13. A. Weymann, E. Hartman, V. Gazit, C. Wang, M. Glauber, Y. Turmelle and D. A. Rudnick, Hepatology, 2009, 50, 207–215.

14. E. P. Newberry, S. M. Kennedy, Y. Xie, J. Luo, S. E. Stanley, C. F. Semenkovich, R. M. Crooke, M. J. Graham and N. O. Davidson, Hepatology, 2008, 48, 1097–1105.

15. E. Shteyer, Y. Liao, L. J. Muglia, P. W. Hruz and D. A. Rudnick, Hepatology, 2004, 40, 1322–1332.

16. W. E. Naugler, PloS one, 2014, 9, e97426.

17. H. H. Otu, K. Naxerova, K. Ho, H. Can, N. Nesbitt, T. A. Libermann and S. J. Karp, The Journal of biological chemistry, 2007, 282, 11197–11204.

18. P. Singh, T. Goode, A. Dean, S. S. Awad and G. J. Darlington, The journals of gerontology. Series A, Biological sciences and medical sciences, 2011, 66, 944–956.

19. M. Pibiri, P. Sulas, V. P. Leoni, A. Perra, M. A. Kowalik, A. Cordella, P. Saggese, G. Nassa and M. Ravo, Age, 2015, 37, 9796.

20. T. A. Zimmers, X. Jin, Z. Zhang, Y. Jiang and L. G. Koniaris, American journal of physiology. Endocrinology and metabolism, 2017, 313, E440–E449.

21. M. P. Valdecantos, L. Ruiz, V. Pardo, L. Castro-Sanchez, C. Garcia-Monzon, B. Lanzon, J. Ruperez, C. Barbas, J. Naylor, J. L. Trevaskis, J. Grimsby, C. M. Rondinone and A. M. Valverde, Scientific reports, 2018, 8, 16461.

22. S. Bangru, W. Arif, J. Seimetz, A. Bhate, J. Chen, E. H. Rashan, R. P. Carstens, S. Anakk and A. Kalsotra, Nature structural & molecular biology, 2018, 25, 928–939.

23. L. Rib, D. Villeneuve, S. Minocha, V. Praz, N. Hernandez, N. Guex, W. Herr and X. C. Cycli, Epigenetics & chromatin, 2018, 11, 52.

24. S. Wang, C. Zhang, D. Hasson, A. Desai, S. SenBanerjee, E. Magnani, C. Ukomadu, A. Lujambio, E. Bernstein and K. C. Sadler, Developmental cell, 2019, 50, 43–56 e46.

25. P. White, J. E. Brestelli, K. H. Kaestner and L. E. Greenbaum, The Journal of biological chemistry, 2005, 280, 3715–3722.

26. M. J. Caldez, M. Bjorklund and P. Kaldis, Hepatology international, 2020, 14, 463–474.

27. K. B. Halpern, R. Shenhav, O. Matcovitch-Natan, B. Toth, D. Lemze, M. Golan, E. E. Massasa, S. Baydatch, S. Landen, A. E. Moor, A. Brandis, A. Giladi, A. S. Avihail, E. David, I. Amit and S. Itzkovitz, Nature, 2017, 542, 352–356.

28. S. Ben-Moshe and S. Itzkovitz, Nat Rev Gastroenterol Hepatol, 2019, 16, 395–410.

29. N. M. Q. Pek, K. J. Liu, M. Nichane and L. T. Ang, Cell Mol Gastroenterol Hepatol, 2021, 11, 273–290.

30. T. Chen, S. Oh, S. Gregory, X. Shen and A. M. Diehl, JCI insight, 2020, 5.

31. U. V. Chembazhi, S. Bangru, M. Hernaez and A. Kalsotra, Genome Res, 2021, 31, 576–591.

32. Y. Wei, Y. G. Wang, Y. Jia, L. Li, J. Yoon, S. Zhang, Z. Wang, Y. Zhang, M. Zhu, T. Sharma, Y. H. Lin, M. H. Hsieh, J. H. Albrecht, P. T. Le, C. J. Rosen, T. Wang and H. Zhu, Science, 2021, 371.

33. L. He, W. Pu, X. Liu, Z. Zhang, M. Han, Y. Li, X. Huang, X. Han, Y. Li, K. Liu, M. Shi, L. Lai, R. Sun, Q. D. Wang, Y. Ji, J. S. Tchorz and B. Zhou, Science, 2021, 371.

34. P. Langfelder and S. Horvath, BMC bioinformatics, 2008, 9, 559.

35. B. Zhang and S. Horvath, Statistical applications in genetics and molecular biology, 2005, 4, Article17.

36. M. V. Kuleshov, M. R. Jones, A. D. Rouillard, N. F. Fernandez, Q. Duan, Z. Wang, S. Koplev, S. L. Jenkins, K. M. Jagodnik, A. Lachmann, M. G. McDermott, C. D. Monteiro, G. W. Gundersen and A. Ma’ayan, Nucleic acids research, 2016, 44, W90–97.

37. M. H. Schulz, W. E. Devanny, A. Gitter, S. Zhong, J. Ernst and Z. Bar-Joseph, BMC systems biology, 2012, 6, 104.

38. E. Mjolsness, D. H. Sharp and J. Reinitz, J Theor Biol, 1991, 152, 429–453.

39. J. J. Tyson and B. Novak, Annu Rev Phys Chem, 2010, 61, 219–240.

40. U. Apte, M. D. Thompson, S. Cui, B. Liu, B. Cieply and S. P. Monga, Hepatology, 2008, 47, 288–295.

41. T. P. Miettinen, H. K. Pessa, M. J. Caldez, T. Fuhrer, M. K. Diril, U. Sauer, P. Kaldis and M. Bjorklund, Current biology : CB, 2014, 24, 598–608.

42. M. Horimoto, P. Fulop, Z. Derdak, J. R. Wands and G. Baffy, Hepatology, 2004, 39, 386–392.

43. Y. Zhao, M. Tran, L. Wang, D. J. Shin and J. Wu, Hepatol Commun, 2020, 4, 504–517.

44. T. P. Miettinen and M. Bjorklund, Trends Cell Biol, 2017, 27, 393–402.

45. H. Zhang, H. Liu, L. Zhou, J. Yuen and H. J. Forman, Free Radic Biol Med, 2017, 113, 304–310.

46. S. C. Lu, Biochimica et biophysica acta, 2013, 1830, 3143–3153.

47. K. J. Riehle, J. Haque, R. S. McMahan, T. J. Kavanagh, N. Fausto and J. S. Campbell, J Liver Disease Transplant, 2013, 1.

48. S. Mukherjee, K. Chellappa, A. Moffitt, J. Ndungu, R. W. Dellinger, J. G. Davis, B. Agarwal and J. A. Baur, Hepatology, 2017, 65, 616–630.

49. S. Hong, J. M. Moreno-Navarrete, X. Wei, Y. Kikukawa, I. Tzameli, D. Prasad, Y. Lee, J. M. Asara, J. M. Fernandez-Real, E. Maratos-Flier and P. Pissios, Nat Med, 2015, 21, 887–894.

50. A. Roberti, A. F. Fernandez and M. F. Fraga, Mol Metab, 2021, 45, 101165.

51. M. Komatsu, T. Kanda, H. Urai, A. Kurokochi, R. Kitahama, S. Shigaki, T. Ono, H. Yukioka, K. Hasegawa, H. Tokuyama, H. Kawabe, S. Wakino and H. Itoh, Scientific reports, 2018, 8, 8637.

52. V. Dubois, C. Gheeraert, W. Vankrunkelsven, J. Dubois-Chevalier, H. Dehondt, M. Bobowski-Gerard, M. Vinod, F. P. Zummo, F. Guiza, M. Ploton, E. Dorchies, L. Pineau Boulinguiez, E. Vallez, E. Woitrain, E. Bauge, F. Lalloyer, C. Duhem, N. Rabhi, R. E. van Kesteren, C. M. Chiang, S. Lancel, H. Duez, J. S. Annicotte, R. Paumelle, I. Vanhorebeek, G. Van den Berghe, B. Staels, P. Lefebvre and J. Eeckhoute, Molecular systems biology, 2020, 16, e9156.

53. A. Klochendler, N. Weinberg-Corem, M. Moran, A. Swisa, N. Pochet, V. Savova, J. Vikesa, Y. Van de Peer, M. Brandeis, A. Regev, F. C. Nielsen, Y. Dor and A. Eden, Developmental cell, 2012, 23, 681–690.

54. H. Wu, T. Reizel, Y. J. Wang, J. L. Lapiro, B. T. Kren, J. Schug, S. Rao, A. Morgan, A. Herman and L. L. Shekels, Proceedings of the National Academy of Sciences, 2020, 117, 17177–17186.

55. E. A. Hanse, D. G. Mashek, J. R. Becker, A. D. Solmonson, L. K. Mullany, M. T. Mashek, H. C. Towle, A. T. Chau and J. H. Albrecht, Cell Cycle, 2012, 11, 2681–2690.

56. F. M. Sladek, Cell Cycle, 2012, 11, 3156–3157.

57. C. Walesky and U. Apte, Gene Expr, 2015, 16, 101–108.

58. F. Garibaldi, C. Cicchini, A. Conigliaro, L. Santangelo, A. M. Cozzolino, G. Grassi, A. Marchetti, M. Tripodi and L. Amicone, Cell death and differentiation, 2012, 19, 937–946.

59. M. K. Jolly, B. T. Preca, S. C. Tripathi, D. Jia, J. T. George, S. M. Hanash, T. Brabletz, M. P. Stemmler, J. Maurer and H. Levine, APL Bioeng, 2018, 2, 031908.

60. A. Mardinoglu, E. Bjornson, C. Zhang, M. Klevstig, S. Soderlund, M. Stahlman, M. Adiels, A. Hakkarainen, N. Lundbom, M. Kilicarslan, B. M. Hallstrom, J. Lundbom, B. Verges, P. H. Barrett, G. F. Watts, M. J. Serlie, J. Nielsen, M. Uhlen, U. Smith, H. U. Marschall, M. R. Taskinen and J. Boren, Molecular systems biology, 2017, 13, 916.

61. M. A. Della Fazia and G. Servillo, Cell Stress, 2018, 2, 162–175.

62. J. Argemi, T. R. Kress, H. C. Y. Chang, R. Ferrero, C. Bertolo, H. Moreno, M. Gonzalez-Aparicio, I. Uriarte, L. Guembe, V. Segura, R. Hernandez-Alcoceba, M. A. Avila, B. Amati, J. Prieto and T. Aragon, Gastroenterology, 2017, 152, 1203–1216 e1215.

63. Y. Liu, M. Shao, Y. Wu, C. Yan, S. Jiang, J. Liu, J. Dai, L. Yang, J. Li, W. Jia, L. Rui and Y. Liu, Journal of hepatology, 2015, 62, 590–598.

64. M. Hamano, H. Ezaki, S. Kiso, K. Furuta, M. Egawa, T. Kizu, N. Chatani, Y. Kamada, Y. Yoshida and T. Takehara, J Gastroenterol, 2014, 49, 305–316.

65. G. Donati, L. Montanaro and M. Derenzini, Cancer research, 2012, 72, 1602–1607.

66. B. Tian and J. L. Manley, Nat Rev Mol Cell Biol, 2017, 18, 18–30.

67. D. Kumar, M. Das, C. Sauceda, L. G. Ellies, K. Kuo, P. Parwal, M. Kaur, L. Jih, G. K. Bandyopadhyay, D. Burton, R. Loomba, O. Osborn and N. J. Webster, The Journal of clinical investigation, 2019, 129, 4477–4491.

68. P. Wu, M. Zhang and N. J. G. Webster, Frontiers in endocrinology, 2021, 12, 613213.

69. C. M. Walesky, K. E. Kolb, C. L. Winston, J. Henderson, B. Kruft, I. Fleming, S. Ko, S. P. Monga, F. Mueller, U. Apte, A. K. Shalek and W. Goessling, Nature communications, 2020, 11, 5785.

70. I. Huck, S. Gunewardena, R. Espanol-Suner, H. Willenbring and U. Apte, Hepatology, 2019, 70, 666–681.

71. N. L. Lazarevich, O. A. Cheremnova, E. V. Varga, D. A. Ovchinnikov, E. I. Kudrjavtseva, O. V. Morozova, D. I. Fleishman, N. V. Engelhardt and S. A. Duncan, Hepatology, 2004, 39, 1038–1047.

72. B. F. Ning, J. Ding, C. Yin, W. Zhong, K. Wu, X. Zeng, W. Yang, Y. X. Chen, J. P. Zhang, X. Zhang, H. Y. Wang and W. F. Xie, Cancer research, 2010, 70, 7640–7651.

73. D. D. Lv, L. Y. Zhou and H. Tang, Exp Mol Med, 2021, 53, 8–18.

74. D. Jia, M. K. Jolly, S. C. Tripathi, P. Den Hollander, B. Huang, M. Lu, M. Celiktas, E. Ramirez-Pena, E. Ben-Jacob, J. N. Onuchic, S. M. Hanash, S. A. Mani and H. Levine, Cancer Converg, 2017, 1, 2.

75. V. Noce, C. Battistelli, A. M. Cozzolino, V. Consalvi, C. Cicchini, R. Strippoli, M. Tripodi, A. Marchetti and L. Amicone, Cell Death Dis, 2019, 10, 768.

76. D. Yimlamai, C. Christodoulou, G. G. Galli, K. Yanger, B. Pepe-Mooney, B. Gurung, K. Shrestha, P. Cahan, B. Z. Stanger and F. D. Camargo, Cell, 2014, 157, 1324–1338.

77. J. L. Grijalva, M. Huizenga, K. Mueller, S. Rodriguez, J. Brazzo, F. Camargo, G. Sadri-Vakili and K. Vakili, Am J Physiol Gastrointest Liver Physiol, 2014, 307, G196–204.

78. O. Croci, S. De Fazio, F. Biagioni, E. Donato, M. Caganova, L. Curti, M. Doni, S. Sberna, D. Aldeghi, C. Biancotto, A. Verrecchia, D. Olivero, B. Amati and S. Campaner, Genes Dev, 2017, 31, 2017–2022.

79. J. O. Russell and S. P. Monga, Annu Rev Pathol, 2018, 13, 351–378.

80. S. P. Monga, Gene Expr, 2014, 16, 51–62.

81. M. Yang, S. N. Li, K. M. Anjum, L. X. Gui, S. S. Zhu, J. Liu, J. K. Chen, Q. F. Liu, G. D. Ye, W. J. Wang, J. F. Wu, W. Y. Cai, G. B. Sun, Y. J. Liu, R. F. Liu, Z. M. Zhang and B. A. Li, J Cell Sci, 2013, 126, 5692–5703.

82. A. Bisso, M. Filipuzzi, G. P. Gamarra Figueroa, G. Brumana, F. Biagioni, M. Doni, G. Ceccotti, N. Tanaskovic, M. J. Morelli, V. Pendino, F. Chiacchiera, D. Pasini, D. Olivero, S. Campaner, A. Sabo and B. Amati, Hepatology, 2020, 72, 1430–1443.

83. L. A. Furchtgott, C. C. Chow and V. Periwal, Biophysical journal, 2009, 96, 3926–3935.

